# Single-cell, single-shot stimulation reveals heterogeneous network recruitment not explained by standard functional properties

**DOI:** 10.64898/2026.06.14.731818

**Authors:** Roy, Offenhäusser, Maybeck

**Affiliations:** Institute of Biological Information Processing (IBI-3), Forschungszentrum Jülich, Germany

**Keywords:** Optogenetics, synchrony, Minimal stimulation, network dynamics, calcium imaging

## Abstract

Understanding how minimal perturbations influence cortical network dynamics remains a central challenge in neural engineering. While single-cell stimulation has been shown to affect population activity, the resulting variability is often treated as noise rather than an informative feature of network behavior. Here, we investigated how single-cell stimulation reflects heterogeneous modulation of neuronal recruitment and the extent to which these effects can be explained by the functional state of the stimulated cell. For this, we combined single-cell optogenetic stimulation with wide-field calcium imaging in cortical cultures. In each network, a single stimulation event was induced, and subsequent alterations in stimulus-coupled recruitment, synchrony, and pairwise correlations were quantified. Additionally, we evaluated whether the baseline functional state of the stimulated neurons, including their event activity levels and Pearson correlation structures, were linked to the observed network responses. Single cell stimulation induced effects were transient, and the network dynamics recovered over a few seconds within the responder population. Importantly, Our findings demonstrate that the observed direction and magnitude of recruitment changes were not significantly explained by the functional state of the stimulated neurons, indicating that these parameters do not capture the determinants of perturbation-induced network responses. This highlights a possibilities in the future approaches for characterizing network responsiveness and suggests that additional unobserved features govern the response of microcircuits to localized inputs.

## 1. Introduction

Modern stimulation strategies increasingly aim to achieve precise and minimally invasive interactions with neural circuits^1^, with applications ranging from basic neuroscience to bioelectronic medicine. However, biological neural networks are inherently dynamic and are rearranged by ongoing activity-dependent plasticity, which results in continuously reconfigured connectivity patterns. This intrinsic variability poses a fundamental challenge: how can minimal localized perturbations influence collective network dynamics in a reliable and interpretable manner?

Although imaging and electrophysiological techniques provide detailed descriptions of evolving network states, they do not establish meaningful relationships between input and network reconfiguration. Perturbation-based approaches address this limitation by introducing controlled inputs and observing the resulting changes in the network activity. Previous studies have demonstrated that even single neurons can influence the local and global circuit dynamics. For example, Kozloski et al. demonstrated that stimulating specific trigger cells within a homogeneous population of corticotectal pyramidal neurons can consistently drive the activity of downstream “follower” neurons^2^. In line with this, a single action potential can initiate complex, temporally extended responses in postsynaptic cells^3^, and the activation of a single pyramidal neuron can evoke reliable sequences of network responses lasting up to approximately 200 ms^4^.

But many of these studies rely on spatial-temporally structured stimulation protocols. While such approaches enhance signal detection and reproducibility, they also introduce history-dependent effects because residual network activity from prior stimuli can influence subsequent responses. This complicates the interpretation of input-output relationships at the level of individual perturbations. At the population level, chronic patterned optogenetic stimulation has been shown to induce functionally distinct cortical ensembles by reshaping intrinsic excitability and synaptic connectivity, leading to reduced correlations between stimulated and non-stimulated populations ^5^. Similarly, brief optical stimulation of a spatially extended area can evoke reliable large-scale network responses ^6^. However, these approaches inherently involve the simultaneous activation of multiple neurons, making it difficult to disentangle the contribution of individual cells to the observed network dynamics. To address this limitation, we adopted a minimal intervention strategy using single-cell, single-shot optogenetic stimulation to isolate the effects of localized inputs on perturbing ongoing network activity. Optogenetic techniques ^7–10^ provide the temporal precision and cellular specificity necessary to test how localized inputs bias the network output. However, it remains uncertain whether responders of such brief single-cell activation can systematically direct the responders in the network toward rearrangement of synchronous network events (SNEs). SNEs capture coordinated participation of neuronal subpopulations and therefore provide a natural framework for assessing how localized perturbations bias network-level organization. For example, Hady et al., 2013, demonstrate that low frequency stimulation protocols can reliably increase burst synchrony, but it’s not known whether a brief pulse can increase the synchrony in between responder populations or stimulation coupled cells within subsequent post stimulation SNEs. Bonifazi et al. demonstrated that specific GABAergic hub neurons can critically shape neuronal synchrony^11^, underscoring how coordinated activity reflects the underlying network organization. Apart from this, connectivity is another fundamental determinant of network behavior.

While structural and morphological connectivity can influence how stimulation propagates through a circuit, neuronal responses are also shaped by indirect interactions mediated by other neurons in the network ^12^. To capture these interactions, functional connectivity can be quantified through correlations in neuronal activity for assessing how neurons dynamically interact during ongoing activity. Importantly, neuronal activity is regulated by homeostatic mechanisms. For instance, Turrigiano et al. showed that synaptic scaling depends on a neuron activity level as part of homeostatic plasticity ^13^. This raises the possibility that targeting neurons with different baseline activity levels may differentially bias network responses, motivating us to investigate whether stimulating highly active neurons compared to less active neurons produces more stable and reproducible network outcomes. For instance, Kobayashi et al. identified leader neurons by characterizing their firing properties, revealing that the bursting activity of individual neurons can initiate network-wide activity ^14^.

In this study, we aimed to characterize the extent to which single-cell and single-shot stimulation can influence network-level activity with spatial and functional specificity. We sought to determine whether minimal perturbations could reliably bias the recruitment of neurons into synchronous network events and whether such responses could be explained by the commonly used functional properties of the stimulated neurons. For this, we quantified how individual perturbations influence neuronal recruitment, synchrony, and correlation structure at the population level. Specifically, we asked the following: (i) how does a minimal perturbation transiently alter the participation of neurons in synchronous activity; (ii) how variable are these effects across stimulation events and cells; and (iii) to what extent can this variability be explained by the functional state of the stimulated neuron.

We found that single cell stimulation elicited transient, network-wide elevations in mean neuronal activity. The responses demonstrated a decay dependent on distance from the stimulated cells, although considerable variability was noted across the networks. These effects persisted for several tens of seconds within the responder population before gradually reverting to baseline levels. Alterations in pairwise correlations were heterogeneous and did not exhibit systematic differences between responder and non-responder cell populations, while the overall structure of synchronous network events remained stable. However, we observed that there is a subtle direction in participation of responders towards the SNEs. Some cell stimulation enriched and others depleted the responders in the following SNEs.

Together, these findings demonstrate that even minimal perturbations can transiently bias network dynamics in diverse ways that are not captured by standard measures of neuronal function such as event activity level and Pearson correlation in between neurons. This highlights the presence of additional unobserved determinants of network responsiveness and underscores the need for new frameworks to characterize how microcircuits transform localized inputs into collective activity. By focusing on single-cell, single-shot perturbations, this study provides a foundation for understanding the limits of controllability in neural systems and informs the development of stimulation strategies that account for intrinsic variability in network dynamics.

## 2. Methods

### 2.1. Neuronal Culture

Cortical neurons were harvested from E18 Wistar rats (RjHan: WI, supplied by Janvier). Neurons were dissociated using 0.05% trypsin for 10 min and plated at a density of 750 cells/mm² onto 70% ethanol-sterilized, poly L-lysine (PLL)-coated Ibidi dishes. Cultures were maintained at 37°C with 5% CO₂ in Neurobasal medium (NB, Thermofisher) supplemented with 0.5 mM L-glutamine (Thermofisher)), 1% B-27 (Thermofisher), and 100 µM gentamycin (Thermofisher). The culture conditions were optimized to promote neuronal growth while minimizing glial proliferation. The medium was half-exchanged twice a week, and experiments began on DIV 21.

### 2.2. Preparation of Poly-L-lysine–Coated Ibidi Dishes

The Ibidi dishes underwent sterilization through washing with 70% ethanol, followed by drying with nitrogen. PLL was diluted at a ratio of 1:100 in Hank’s Balanced Salt Solution (HBSS) to achieve a final concentration of 10 µg/mL. Approximately 500 µL of the PLL solution was then added to each dish to ensure complete coverage of the bottom surface. The dishes were incubated at room temperature for 60 minutes, after which they were washed at least twice with HBSS at room temperature to eliminate excess PLL. Subsequently, the dishes were sealed with Parafilm and stored at 4 °C for a maximum of two weeks prior to use.

### 2.3. Transduction

Primary neuronal cultures were double-transduced with adeno-associated virus (AAV). Viruses were added to the supplemented NB medium on DIV 7 in adequate dilution, leading to a multiplicity of infection (MOI) of 4.395E5 genome copies/cell (GC/cell) for AAV9 pAAV.Syn.NES-jRCaMP1b. WPRE.SV40 (Plasmid #100851 was obtained from Addgene) ^15^; 0.25E5 GC/cell for AAV8 pAAV-Syn-ChR2(H134R)-GFP (Plasmid #58880 was obtained from Addgene)^7^. At the next bi-weekly medium change (3-5 days later), the virus-containing medium was completely exchanged with supplemented NB medium that did not contain AAVs to remove excess virus particles that did not infect the cells. The transduction efficiency was >80%. Transduction efficiency was quantified as the fraction of cells expressing RCaMP (RFP fluorescence) under an EVOs microscope (Axiovert 200; Zeiss, Oberkochen, Germany) relative to the total number of cells per field of view, where the total number of cells was identified by bright-field-based cell counting. To reduce underestimation due to intermittent RCaMP fluorescence, each field was recorded as a short time-lapse, and cells were classified as RFP-positive if fluorescence exceeded the threshold in any frame.

### 2.4. Calcium Imaging and LASER stimulation

Calcium imaging was performed using an Axio Observer Z1/7 microscope (Zeiss) equipped with an EpiPlan 10x/0.45 objective. The imaging acquisition parameters included a pixel scaling of 1.3 µm × 1.3 µm, an image resolution of 1280 × 1080 pixels, and a 16-bit depth. Calcium imaging was performed using a green LED light source (Zeiss) with 50% intensity, an excitation wavelength of 555 nm; emission was filtered through a custom-made filter (LP590-640 nm, BS580). The acquisition was performed with an exposure time of 20 ms, 2 × 2 binning, and a frame rate of approximately 28 frames per second.

Optogenetic stimulation experiments were performed using a Rapp Optoelectronic system equipped with a 473 nm blue laser, which provided precise spatial and temporal control of light delivery. Laser stimulation was confined to a defined area of approximately 314.16 µm², estimated from the full width at half maximum (FWHM) of the laser intensity profile. corresponding to a single neuronal soma. A single pulse of 100 ms duration was delivered per trial. Stimulation was applied at 5% of the maximum laser power (110 mW source power), corresponding to an estimated output of 0.48 mW at the sample plane, ensuring reliable activation of ChR2-expressing neurons while minimizing the off-target effects.

Calcium-dependent fluorescence signals were recorded before, during, and after stimulation and used as readouts of neuronal and network activity. Imaging data were preprocessed using Suite2p for region of interest (ROI) extraction, and fluorescence trace generation^16^. Subsequent analyses were performed using custom Python scripts employing NumPy, SciPy, and Matplotlib for event detection, quantification, and visualization.

### 2.5. Data Analysis Methods

#### 2.5.1. Data Preprocessing and ΔF/F Computation

Raw fluorescence traces were extracted using Suite2p with standard parameters (threshold = 0.2, S/N ratio: 0.5). Valid cells were identified using the built-in classifier (iscell.npy). For each valid cell, the ΔF/F signal was computed as:

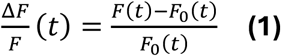

where *F*_0_(*t*) is a sliding percentile baseline (5th percentile, 60-second window) computed using a frame-centered sliding window approach. Stimulus artifact frames (stim_frame − 3 to stim_frame) were set to NaN before baseline computation.

#### 2.5.2. Calcium Event Detection

Discrete calcium events were detected from ΔF/F traces using a sliding baseline z-score threshold approach. For each cell, a local baseline mean *μ*(*t*) and standard deviation *σ*(*t*) were estimated within a centered sliding window (window size = 60 s). A binary event was registered when:

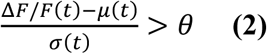

where *θ* = 2 SD. Detected events were required to be separated by a minimum inter-event interval of 20 frames (∼0.69 s at 28.88 Hz) to avoid double-counting of the same calcium transient.

#### 2.5.3. Responder Cell Classification

Responder cells were defined as cells exhibiting at least one calcium event within a post-stimulation detection window of 20 frames (∼0.69 s) immediately following the stimulus onset frame. The stimulated cell itself was excluded from responder classification. For each dataset, the responder percentage was computed as:

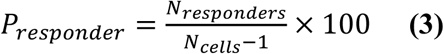

Cell positions were extracted from Suite2p stat.npy using the med field (centroid coordinates), scaled by a pixel size of 1.3 μm/pixel. Euclidean distances from the stimulated cell were computed for each responder and binned into spatial distances.

Spatial response probability was computed per distance bin as the fraction of cells that responded to stimulation among all cells within that bin. Probability distributions were compared across biological replicates using non-parametric statistics (Kruskal–Wallis followed by Dunn’s post-hoc test with Bonferroni correction).

#### 2.5.4. Responder vs. Non-Responder Comparison

For each dataset, two network metrics were compared between responder and non-responder populations:

Pairwise functional correlation: mean Pearson correlation coefficient between all cell pairs within a group, computed after Fisher z-transformation to reduce averaging bias:

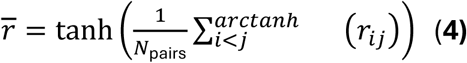

Population activity level: mean event rate (events per second) averaged across group members. Group differences were assessed using the Wilcoxon signed-rank test across N = 89 networks.

#### 2.5.5. Stimulated Cell Characterization and Classification

The stimulated cell in each dataset was characterized along two independent axes derived from the 60-second pre-stimulation baseline:

- Baseline correlation: mean pairwise Pearson correlation of the stimulated cell with all other valid cells.
- Baseline activity: mean calcium event rate (events per second).

Cells were selected randomly during the experiment and post-hoc categorized into one of four types using the median splits for correlation and activity to define high and low performing cells for each metric in a given network. We have chosen a random single cell from each network and then those cells were classified into four categories using median splits of both metrics in each network:

Cells were classified into four categories based on median splits of correlation and activity within each network. Correlation values below a threshold of 0.2 were considered low regardless of the median. Accordingly, cells were categorized as high correlation / high activity (correlation ≥ median and ≥ 0.2; activity ≥ median), high correlation / low activity (correlation ≥ median and ≥ 0.2; activity < median), low correlation / high activity (correlation < median or < 0.2; activity ≥ median), and low correlation / low activity (correlation < median or < 0.2; activity < median). Category membership was tested for association with post-stimulation network enrichment using a chi-squared test of independence.

#### 2.5.6 SNE Analysis

To detect SNEs, population activity was quantified using event onsets. We derived the onset timing (o) from the event matrix (e). For each cell i, an onset indicator *o*_*i*_(*t*) ∈ {0,1} was defined such that *o*_*i*_(*t*) = 1 when an event began at frame **t** *e*_*i*_(*t*) = 1⍰ *e*_*i*_(*t* − 1) = 0⍰)*o*_*i*_(*t*) = 0⍰ otherwise. Population onset activity was then computed frame-by-frame as the fraction of cells whose events started at that frame:

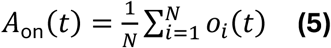

Where **N** is the total number of valid cells in the recording.

An SNE was identified when population onset activity exceeded a dynamic threshold defined as:

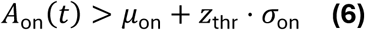

where *μ*_on_ and *σ*_on_ represent the mean and standard deviation of *A*_on_(*t*) computed across the full recording, and *z*_thr_ = 2.0. This z-score threshold was selected to detect statistically significant population-level synchrony while minimizing detection of stochastic co-activation of small cell subsets.

For each detected SNE, the participating cell ensemble was defined as cells *e*_*i*_(*t*) = 1⍰ in at least one frame within the SNE window[*t*_onset_, *t*_offset_] 692 ms ⍰ Participation rate for cell **i** across all SNEs was computed as:

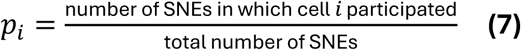

Cells with *p*_*i*_ > 0.80 were classified as core SNE members (high-fidelity participants), and those with *p*_*i*_ ≤ 0.80 as peripheral members.

To test whether optogenetically recruited responder cells preferentially participate in subsequent SNEs, the overlap between the responder ensemble and the post-stimulation SNE core membership was quantified using the Jaccard similarity index:

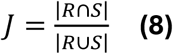

Where **R** is the set of responder cells and **S** is the set of post-stimulation core SNE members.

For each SNE, the fraction of responder cells among its participants was computed as:

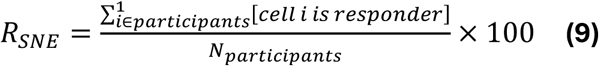

The mean responder fraction was then averaged across all baseline SNEs and all post-stimulation SNEs separately:

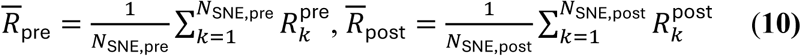

Enrichment was defined as the absolute difference:

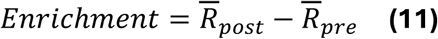

## 2. Results

### 3.1. Single-cell optogenetic stimulation (SCOC) reliably elicits cellular responses and evokes transient network-level activity

To explore the influence of a single neuronal stimulation on population dynamics, we conducted calcium imaging in neuronal cultures expressing optogenetic actuators and genetically encoded calcium indicators (Fig. 1A). Fluorescence imaging confirmed robust expression throughout the network, facilitating the simultaneous monitoring of activity in extensive neuronal populations. Initially, we validated the efficacy of single-cell stimulation at the individual neuron level. Targeted optogenetic activation induced a rapid and transient increase in calcium fluorescence (ΔF/F), demonstrating reliable activation of stimulated neurons (Fig. 1B). Across experiments, stimulation recruited a variable fraction of neurons within the network, with responder percentages exhibiting a median of 12.36% and spanning a broad range (Fig. 1C), indicating substantial heterogeneity in network responsiveness. To evaluate the impact of stimulation on global network dynamics, we constructed raster plots of the inferred activity across all recorded neurons (Fig. 1D). Spontaneous activity was characterized by intermittent and synchronous population events. Alignment of activity to stimulation onset revealed that single-cell activation was followed by coordinated responses across subsets of neurons (responders) within a short temporal window, suggesting that perturbation of a single neuron biases ongoing network activity. Notably, the extent of recruitment varied across trials, supporting the notion that single-cell stimulation does not deterministically trigger network events but instead increases the probability of a population-wide activation.

**Figure 1:**
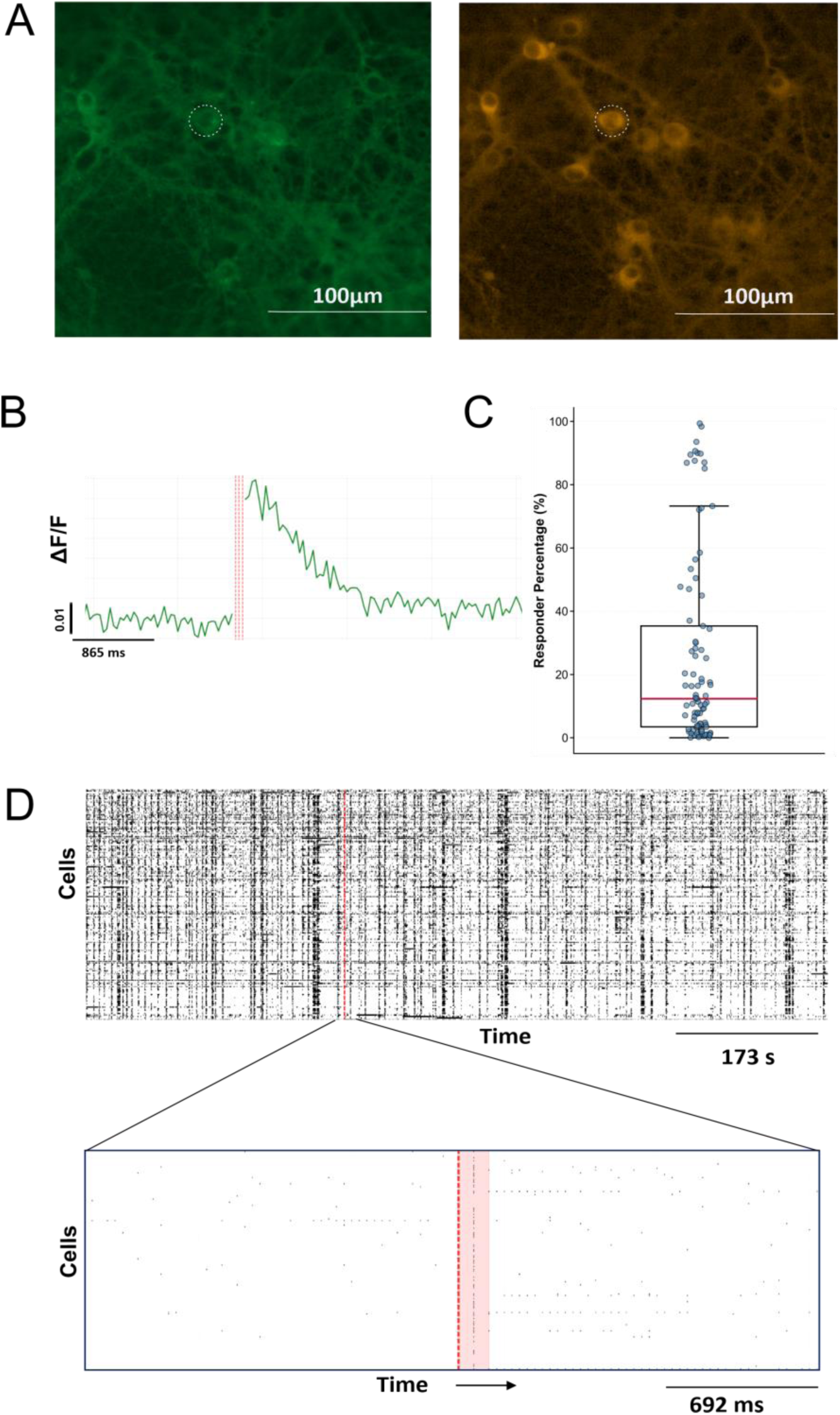
A) Co-expression of optogenetic actuator ChR2-GFP (left) and calcium indicator RCaMP (right) in neuronal culture. B) SCOC reliably evokes a response to the targeted cell. Normalized fluorescence signal (dF/F) by 5^th^ percentile method. C) The response is not restricted to a target cell; a variable percentage of responders in the network respond to stimulation, Median: 12.36%, Interquartile Range (IQR, 25^th^-75^th^): 3.48% - 35.36%. D) Network-wide activity patterns and temporally aligned responses to single cell stimulation (top), red line = stimulation. Bottom: Zoom view of stimulus coupled zone.

To determine whether stimulation-induced responses exhibit spatial organization, we initially analyzed the spatial distribution of responders and non-responders across the network (Fig. 2A). Although responsive neurons were frequently located near the stimulated cells, numerous non-responders were also observed in close proximity, indicating a considerable local variability. We quantified this relationship by evaluating the probability of neuronal activation as a function of the distance from the stimulated cell (Fig. 2B). Spatial mapping revealed that the recruited cells were distributed throughout the network, with a higher density of responders near the stimulation site. Consistent with this observation, the response probability was highest within the nearest distance bin (0–100 µm) and decreased progressively with increasing distance. Quantitative analysis across all networks confirmed a significant distance-dependent decline in response probability, with proximal neurons exhibiting a higher likelihood of activation than more distal populations. However, responses at greater distances, although less frequent, remained detectable and more variable. Collectively, these findings indicate that single-cell stimulation exerts a spatially graded influence, preferentially recruiting nearby neurons while still affecting the distributed network elements.

**Figure 2:**
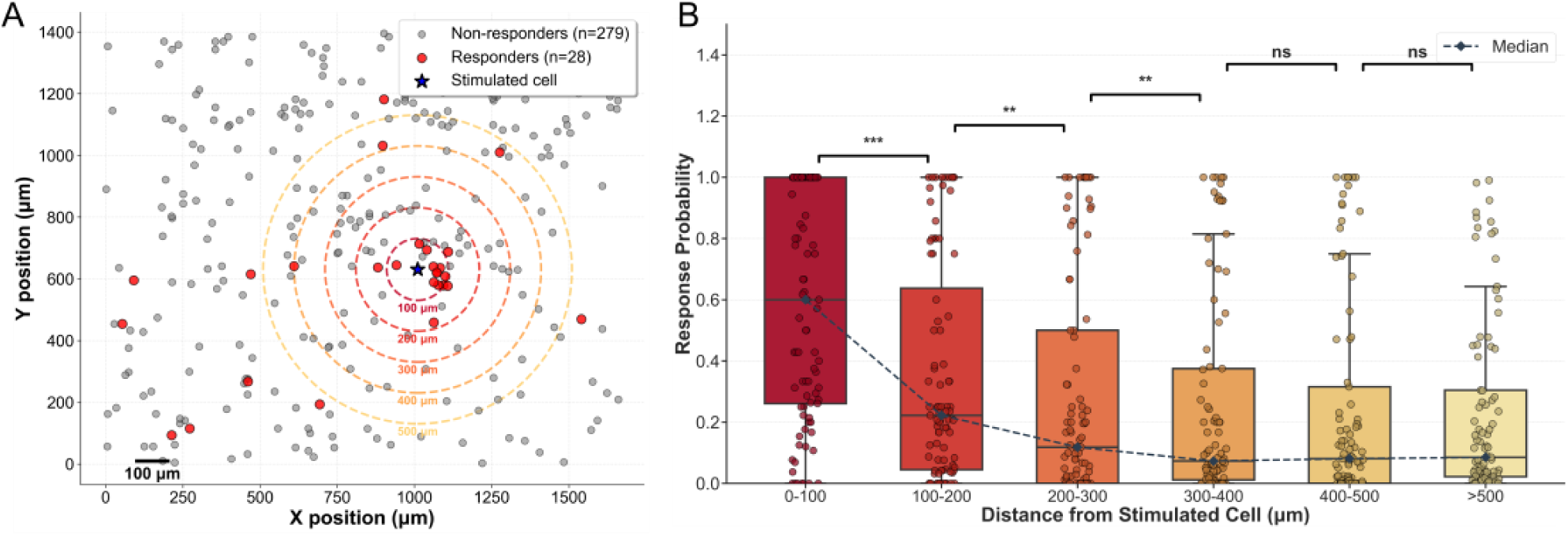
Spatial distance alone does not account for variability in neuronal recruitment. A) Spatial representation of all active cells in the network. Cells with a calcium event within 700 ms of stimulation (responders) are marked red. The stimulated cell is marked by the blue star. Concentric rings are a guide to the eye of distance from stimulated cell showing how cells were classified for results shown in B. B) Response probability decreased significantly up to 400 μm, afterwards the percent of responders becomes stable [*** = p < 0.001, ** = 0.001< p < 0.01, * = 0.01< p < 0.05 ns = p ≥ 0.05]

### 3.2 ) Responder-specific modulation does not alter the ensemble structure of network activity

Responder neurons are characterized by calcium activity that is temporally aligned with the optogenetic pulse, indicating a stimulation-coupled subpopulation within the network. To determine whether this coupling signifies functional differences in network engagement, we compared responders to non-responder neurons regarding their overall activity and pairwise correlations. Following stimulation, both responders (***p < 0.001) and non-responders (**p < 0.01) demonstrated a significant increase in mean activity post-stimulation. This suggests that stimulation preferentially enhances activity within a defined subpopulation rather than uniformly across the network (Fig. 3A). This selectivity was even more pronounced at the level of network coordination. The pairwise correlations among responders were significantly altered following stimulation (*p < 0.05), indicating a reorganization of coordinated activity within this group. In contrast, non-responders showed no significant change in the correlation structure, suggesting that stimulation-driven network reconfiguration is largely confined to the responder’s subnetwork (Fig. 3B).

**Figure 3:**
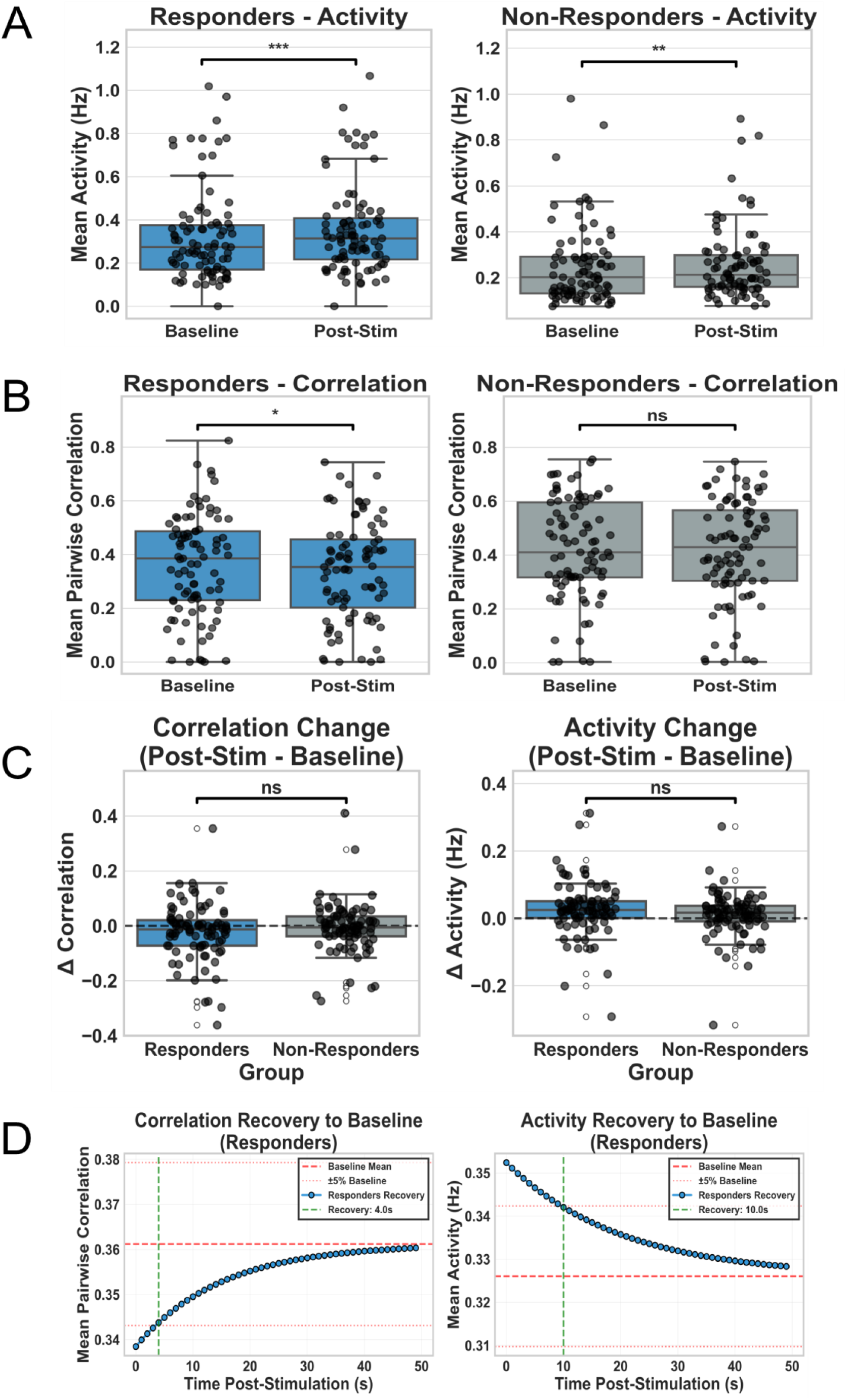
Activity and functional coupling in responder and non-responder neurons before and after stimulation. (A) Mean neuronal activity (Hz) for responders and non-responders at baseline and following stimulation. (B) Mean pairwise correlation for responders and non-responders before and after stimulation. (C) Changes in activity and correlation (post-stimulation minus baseline) comparing responder and non-responder populations. (D) Temporal evolution of correlation and activity following stimulation in responders, demonstrating recovery toward baseline levels over time.

Notably, these within-group effects were not reflected in direct comparisons between cell types. The change of both activity and correlation did not differ significantly between the responders and non-responders (Fig. 3C). Thus, despite apparent differences in within-group responses, cell-type-specific effects cannot be distinguished at the level of global summary metrics. However, these effects were transient, as both activity and correlation in responders gradually returned to baseline levels over time; specifically, transient correlation returned to baseline at 4 seconds, while activity recovered after 10 seconds (Fig. 3D).

The simultaneous activation of multiple responders following single cell stimulation appears to resemble SNEs, prompting us to quantitatively investigate their contribution to spontaneous SNEs. Specifically, we aimed to evaluate whether externally induced activity is assimilated into ongoing network dynamics through the increased involvement of responders in SNEs. The similarity of the SNEs, as quantified by the Jaccard index, revealed overlapping distributions between the baseline and baseline–post-stimulation comparisons (Fig. 4A), with most values ranging from approximately 0.7 to 1.0. This finding suggests that the composition of co-active neuronal groups is largely preserved after stimulation. Importantly, the lack of a pronounced shift or clustering within the distribution indicates that stimulation does not systematically reorganize SNE membership but rather maintains the existing ensemble structure with only minor variability across the events. Consistent with this, the fraction of responders participating in individual SNEs exhibited no systematic change following stimulation (Fig. 4B).

**Figure 4:**
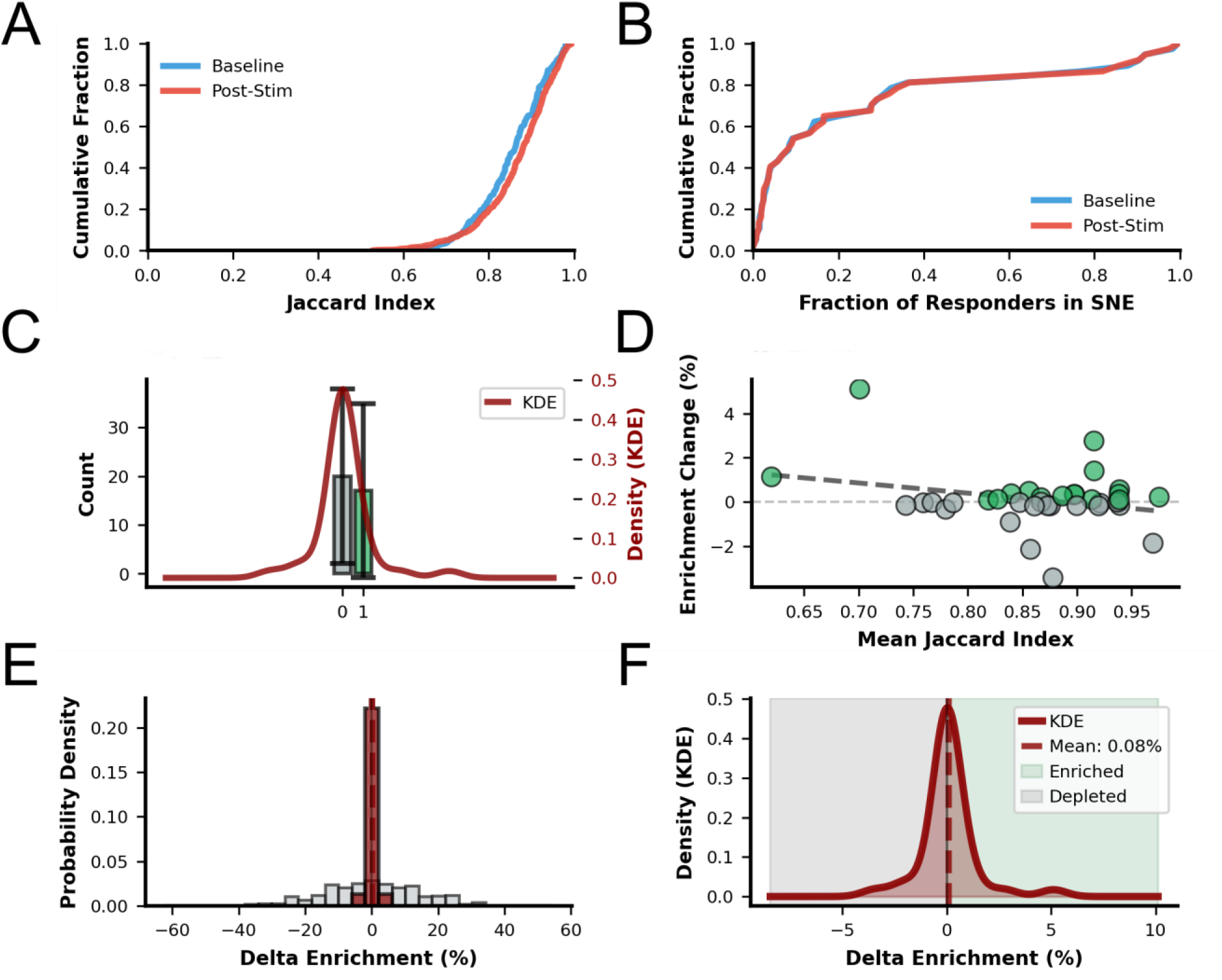
(A) Comparison of baseline (blue) and post-stimulation (red) SNE assembly similarity, quantified as the Jaccard index of cell-membership overlap between consecutive SNE pairs, showing no global shift in network structure (Wilcoxon signed-rank test: W = 113.0, p = 0.1165, rank-biserial r = 0.356; median Jaccard: baseline = 0.844 [IQR 0.812–0.891], post = 0.876 [IQR 0.849–0.917]). (B) Cumulative distribution of responder participation in SNEs before (blue) and after (red) stimulation, indicating no consistent increase in recruitment (Wilcoxon signed-rank test: W = 317.0, p = 0.6120, r = 0.098; median responder fraction: baseline = 0.085 [IQR 0.026–0.307], post = 0.090 [IQR 0.024–0.319]). **(C)** Distribution of enrichment relative to a parametric shuffle-control null model (10 permutations per session, 370 null samples total), demonstrating that observed changes fall within expected spontaneous variability. Sessions showing (green box - 1) enrichment: n = 17/37; (gray box - 0) depleted: n = 20/37. Two-sided binomial test (H₀: P(enriched) = 0.5): p = 0.7428, observed proportion = 0.459 (95% CI 0.289–0.638). The overlaid KDE (dark red, right axis) shows the continuous enrichment value distribution. **(D)** Relationship between each session’s mean post-stimulation Jaccard index and its enrichment change, showing no significant association (Pearson r = −0.275, R² = 0.076, Cohen’s f² = 0.082 [small], p = 0.1045; *n* = 37). Points colored green (enriched, n = 17) or grey (depleted, n = 20); dashed line = linear regression fit. (E) Distribution of stimulation-induced enrichment changes across sessions. Real delta enrichment (dark red; mean = 0.085%, SD = 1.284%) compared against the shuffle-control null distribution (light grey; mean = −0.035%, SD = 15.266%; 370 null samples). Despite the real distribution being significantly narrower than the null (two-sample KS test: D = 0.419, p < 0.001), the real enrichment values themselves do not deviate from zero. Dashed vertical lines mark respective means; solid black line = zero. (F) Kernel density estimate of enrichment shifts across sessions (dark red), centered near zero and exhibiting bidirectional variability (mean = 0.085%, SD = 1.284%; one-sample Wilcoxon signed-rank test vs 0: W = 317.0, p = 0.6120, rank-biserial r = 0.098, Cohen’s d = 0.066 [small]). Dashed line = mean enrichment; background shading indicates enriched (x > 0, dark green) and depleted (x < 0, dark grey) regions.

To determine whether stimulation led to preferential incorporation of responders into coordinated activity, we compared enrichment against a null model. We found that enrichment values were not significantly different from null expectations (Fig. 4C, 4E, 4F), indicating that observed variations in responder participation fall within the range of spontaneous network fluctuations. Furthermore, variability in enrichment across recordings was not significantly associated with ensemble similarity (Fig. 4D), suggesting that these fluctuations are not structured by the underlying network organization. Together, these results indicate that, although single-cell stimulation is associated with variability at the level of individual neuronal responses, it does not produce detectable reorganization of coordinated network activity. Instead, synchronous network events remain stable, and changes in responder participation are consistent with spontaneous variability.

### 3.3 ) Network-coupling properties of the stimulated neuron

Given that stimulation-induced effects do not reorganize the structure of SNEs, we next investigated whether the functional state of the stimulated neuron influences the interaction of stimulation with this stable network architecture. Specifically, we tested whether the intrinsic properties of the stimulated cell, such as its baseline activity and coupling to the network, were associated with differences in recruitment patterns. Analysis of randomly chosen stimulated neurons revealed substantial variability in both mean activity and pairwise correlation with the surrounding population (Fig. 5A–5B), indicating that the neurons occupy diverse functional states within the network. These properties define a spectrum ranging from highly active and strongly coupled neurons to cells that are either weakly active or weakly integrated into the neural network. To capture this heterogeneity, the stimulated neurons were categorized based on their combined activity and correlation profiles. This classification revealed distinct subpopulations with differing degrees of network coupling and intrinsic excitability (Fig. 5D). Importantly, these differences do not reflect changes in the ensemble structure but rather variations in the positioning of individual neurons within that structure.

**Figure 5:**
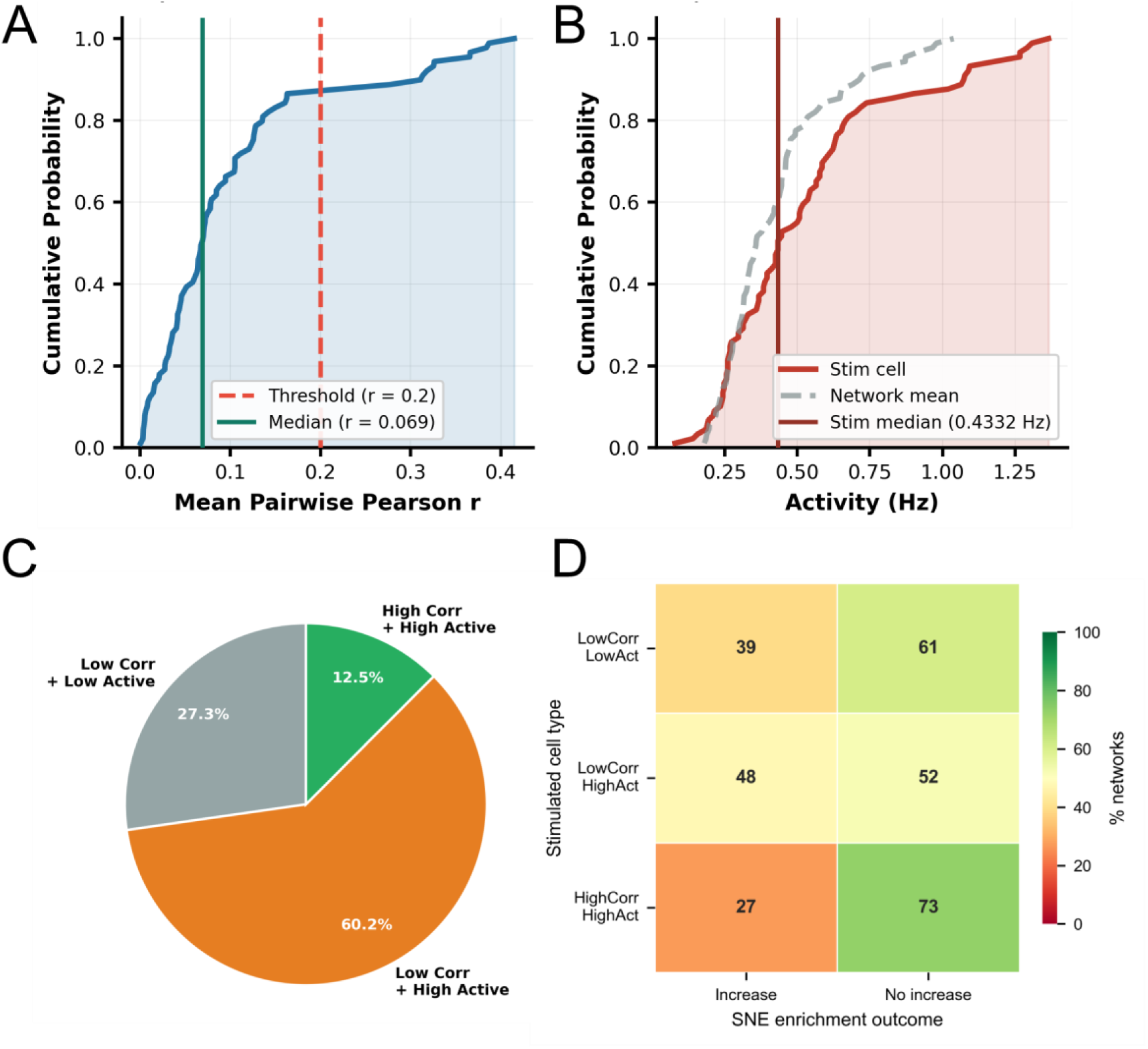
Characterization of stimulated neurons based on baseline activity and functional coupling. (A) CDF of the mean pairwise Pearson correlation between each stimulated cell and all other network cells during baseline. Red dashed line = classification threshold (r = 0.20); green solid line = median across networks. (B) CDF of calcium event onset frequency (Hz) for stimulated cells (red) and per network mean (grey dashed line). Dark red vertical line = median stimulated cell activity. A cell is classified as highly active if its rate exceeds the session network mean. (C) Proportion of networks grouped by the combined baseline profile of the stimulated cell: Low Corr + Low Active (grey), Low Corr + High Active (orange), High Corr + High Active (green). Classification thresholds: r ≥ 0.20 for correlation; activity above network mean. No formal statistical tests are shown in this figure. (D) Enrichment outcome across stimulated cell functional classes. Heatmap shows the percentage of networks exhibiting an increase or no increase in SNE enrichment for each class of stimulated neurons, categorized based on baseline activity and functional correlation (see Methods for threshold definitions).

To ascertain whether responder enrichment was contingent upon the functional state of the stimulated neuron, the preparations were categorized based on the activity of the stimulated cell and network correlation. Across all categories, enrichment outcomes exhibited a broad distribution, and no significant association was identified between the type of stimulated cell and the direction of enrichment (χ² = 1.76, p = 0.415; Figure 6). Although numerical differences in binder fractions were evident across groups, they did not achieve statistical significance, indicating that the likelihood of enrichment was not predicted by the activity or correlation or combination thereof in the stimulated cell.

We observed that randomly sampled neurons did not include cells exhibiting both high functional correlation and low baseline activity. This indicates that, within the recorded networks, strongly correlated neurons tend to also exhibit elevated activity levels, reflecting an intrinsic coupling between participation in network dynamics and baseline excitability. This suggests that functional connectivity and baseline activity are not independent features in these networks, consistent with prior observations that highly connected neurons often occupy central and active roles in coordinating network dynamics^11^.

## 3. Discussion

In neural engineering, it remains challenging to consistently influence collective neural dynamics with minimal intervention. In this study, we quantitatively demonstrated how even a single calcium event can transiently influence network-level activity; however, these effects are highly heterogeneous and bidirectional. Rather than producing consistent or uniform outcomes, individual stimulation events led to bidirectional outcomes, highlighting that minimal perturbations can bias ongoing network dynamics without exerting deterministic control. These observations emphasize the sensitivity of neuronal microcircuits to localized input, while underscoring the limits of predictability at this scale.

We focused on SNEs as a readout of network response because they capture coordinated population activities rather than isolated neuronal responses ^17^. This makes them particularly suitable for assessing the influence of localized perturbations on collective dynamics. Homeostatic plasticity mechanisms have been shown to regulate neuronal excitability in an activity-dependent manner ^18^, suggesting that highly active neurons may occupy privileged positions in maintaining network stability ^13^. Conversely, neurons that are weakly correlated with ongoing network activity are less likely to be embedded within recurrently co-active assemblies and may therefore be expected to exert a weaker influence on coordinated population responses, consistent with Hebbian principles ^19^. A key finding of this study is that the direction of responder enrichment and depletion in network responses to single-cell stimulation cannot be explained by commonly used functional descriptors. Specifically, baseline firing activity and Pearson correlation did not show a significant association with the direction of the stimulation-induced changes. This challenges the assumption that functional embedding, as captured by standard measures such as activity level or pairwise correlation, is sufficient to determine how neurons influence network dynamics.

Previous work suggests that cortical microcircuits can exhibit a measurable sensitivity to perturbations at the level of single neurons, for example, A. C. Kwan and Yang Dan demonstrated that stimulating individual neurons can influence local network activity in vivo^20^, while Michael London and colleagues showed that cortical networks can also operate in a regime of high variability^21^, where responses to small perturbations are inconsistent and noise sensitive. In this study, we probed microcircuit sensitivity by applying controlled single-cell perturbations and quantifying their impact on coordinated population activity. But our studies had several limitations. First, we focused on single-shot stimulation and did not assess the reproducibility of the responses across repeated perturbations of the same neuron. This was a deliberate design choice, motivated by evidence that burst-like perturbations can leave transient traces in network activity that bias subsequent responses^22,23^. Second, our analysis relies on functional connectivity inferred from activity correlations, which captures statistical dependencies but does not provide direct information about the underlying synaptic structure. Consequently, future work should investigate underlying synaptic architecture, including connectivity, synaptic strengths, and recurrent motifs. Third, our experiments were performed in vitro. The core finding that single-neuron perturbations can recruit coordinated and variable population responses is likely to generalize local cortical microcircuits. And it remains to be determined how these findings generalize in vivo circuits with more complex dynamics and modulatory influences on the brain. Fourth, the variability observed in network responses may also reflect differences in the identity of the stimulated neurons. In vitro, cortical cultures contain a heterogeneous mixture of excitatory and inhibitory neurons, yet stimulation targets in this study were not selected based on cell-type-specific markers. As morphological features alone are insufficient for reliable classification, we cannot determine whether the stimulated neurons were excitatory or inhibitory. Kwan et al. reported that the stimulation of a single neuron can elicit variable spatial responses depending on whether the targeted neuron is excitatory or inhibitory ^24^. Therefore, future studies should test whether cell-type identity contributes to the observed variability by repeating the experiments using cell-type-specific stimulation of excitatory and inhibitory neurons.

Despite these limitations, our results have important implications for neural engineering. Many stimulation strategies implicitly assume that increasing stimulation strength23,^26^ or targeting larger populations^4,27^ will yield more reliable response over network activity. In parallel, recent advances in optical stimulation technologies have enabled increasingly precise manipulation of neural circuits at cellular resolution. For example, John P. Rickgauer, Karl Deisseroth and David W. Tank demonstrated that targeted stimulation of individual neurons can influence the activity of small, functionally related populations in vivo, while Adam M. Packer and colleagues developed all-optical approaches for manipulating and recording activity with single-cell and ensemble-level precision. These approaches reflect an ongoing effort to achieve increasingly specific and controlled perturbations of neural circuits^28,29^. In contrast, our findings reveal that single-cell stimulation induces transient modulation of neuronal activity and variable specificity in neuronal recruitment; however, the responses remain unpredictable. We observed an increase in mean activity from approximately 0.30 to 0.40 Hz among responders, while non-responders exhibited activity levels of approximately 0.20–0.25 Hz (p < 0.01). This indicates that perturbation of a single neuron can influence the distributed network activity beyond the directly responsive cells within the microcircuit. These effects persisted for several tens of seconds before gradually returning to baseline levels, suggesting that even minimal perturbations can leave measurable but reversible traces in the ongoing network dynamics. Notably, the probability of response decreased with increasing distance from the stimulated cell (from approximately 0.6 in the 0–100 µm range to less than 0.2 beyond 400 µm); however, substantial variability persisted within each distance range. This suggests that while spatial proximity constrains the spread of influence, it is insufficient to fully predict recruitment in the microcircuits. Furthermore, changes in pairwise correlations were heterogeneous and not significantly different between responders and non-responders, and the structure of the synchronous network events remained stable. This implies that stimulation does not globally reorganize the network structure but modulates the participation of neurons within an otherwise stable dynamic framework. However, the direction of this influence, whether stimulation increases or decreases the participation of responders in synchronous network events, cannot be determined from the baseline event rate and Pearson correlation of stimulated neurons. These findings suggest that improving stimulation specificity may require not only precise control of stimulation parameters but also a deeper understanding of the latent structure and state dependence of neural network. The observed variability, together with the lack of explanation by standard functional descriptors, indicates that the underlying principles governing this influence remain unknown. Identifying features that better capture network responsiveness can inform the design of adaptive stimulation strategies.

In summary, these findings highlight a gap in our current understanding of how microcircuits respond to minimal perturbations and suggest that additional unobserved factors govern network responsiveness. By framing variability as an informative feature rather than noise, this study provides a foundation for developing more nuanced models of neural control and designing stimulation strategies that account for the intrinsic complexity of neuronal networks. One approach to future stimulation designs could be to integrate multi-site, context-aware perturbation strategies that investigate and leverage local microcircuit structures. This includes the sequential or combinatorial activation of neurons to infer and guide network responses beyond the capabilities of isolated, single-cell targeting.

## 4. Conclusion

In conclusion, this study demonstrates that brief single-cell optogenetic stimulation can influence collective network dynamics by biasing the probability of neuronal participation in synchronous network events. These effects are heterogeneous and bidirectional, indicating that minimal perturbations can transiently reshape ongoing coordination patterns without exerting a deterministic control. Importantly, the direction of this influence, whether participation increases or decreases, cannot be explained by the commonly used functional properties of the stimulated neuron (activity and correlation), highlighting a limitation in the current descriptors of network organization. Rather than reflecting random variability, these responses reveal a structured yet unresolved aspect of network dynamics in which single-cell perturbations alter the probability landscape of collective activity. Together, these findings suggest that effective modulation of neural networks will require approaches that account for intrinsic variability and the currently uncharacterized factors governing network responsiveness, providing a foundation for the future development of adaptive and state-aware stimulation strategies.

## Acknowledgements

This research was funded by the Helmholtz Gemeinschaft. We express our gratitude to Bettina Breuer for her exceptional assistance with cell culture. ChatGPT by OpenAI and GitHub Copilot were used only to assist with language refinement and Python code development/debugging; all outputs, analyses, interpretations, and final text were reviewed and verified by the authors.

## Data availability statement

The data that support the findings of this study are available from the authors upon reasonable request.

## Ethics statement

Primary neurons were isolated in compliance with the German Animal Protection Law (TSchG). All experimental protocols received approval from the local animal ethics committee (Landesumweltamt für Natur, Umwelt und Verbraucherschutz (LANUV), Nordrhein-Westfalen, Recklinghausen, Germany; #number 81-02.04.2023.A172). All methods were conducted in accordance with relevant guidelines and regulations and are reported in compliance with ARRIVE guidelines.

## Author contributions

S.R. contributed to conceptualization, experimental design, data collection, data analysis, and manuscript authorship. V. M. advised conceptualization, experimental design, result interpretation, and manuscript composition. A.O. provided the funding for the work and contributed feedback on the manuscript.

